# We see what we hear: dissonant music engages early visual processing

**DOI:** 10.1101/2023.07.07.548089

**Authors:** Fernando Bravo, Jana Glogowski, Emmanuel Andreas Stamatakis, Kristina Herfert

## Abstract

The neuroscientific examination of music processing in audiovisual contexts offers a valuable framework to assess how auditory information influences the emotional encoding of visual information. Using fMRI during naturalistic film viewing, we investigated the neural mechanisms underlying music’s effect on valence inferences during mental state attribution. Thirty-eight participants watched the same short-film accompanied by systematically controlled consonant or dissonant music. Subjects were instructed to think about the main character’s intentions. The results revealed that increasing levels of dissonance led to more negatively-valenced inferences, displaying the profound emotional impact of musical dissonance. Crucially, at the neuroscientific level and despite music being the sole manipulation, dissonance evoked the response of the primary visual cortex response (V1). Functional/effective connectivity analysis showed a stronger coupling between the auditory ventral stream (AVS) and V1 in response to tonal dissonance, and demonstrated the modulation of early visual processing via top-down feedback inputs from the AVS to V1. These V1 signal changes indicate the influence of high-level contextual representations associated with tonal dissonance on early visual cortices, serving to facilitate the emotional interpretation of visual information. The findings substantiate the critical role of audio-visual integration in shaping higher-order functions such as social cognition.

**Significance statement:** The present study reveals responses in the primary visual cortex modulated by musical information: tonal dissonance recruits early visual processing via feedback interactions from the auditory ventral pathway to the primary visual cortex. We demonstrate that the auditory “what” ventral stream plays a role in assigning meaning to non-verbal sound cues, such as dissonant music conveying negative emotions, providing an interpretative framework that serves to process the audio-visual experience. Our results highlight the significance of employing systematically controlled music, which can isolate emotional valence from the arousal dimension, to elucidate the brain’s sound-to-meaning interface and its distributive crossmodal effects on early visual encoding during naturalistic film viewing.

**Data sharing:** All relevant data are available from the figshare database DOI: 10.6084/m9.figshare.21345240

## Introduction

We share with our ancestors the ability to identify other’s intentions based on the available sensory information. The concept of neocortical operations being multisensory is widely accepted and emphasizes the importance of integration in sensory encoding (Ghazanfar & Schroeder, 2006). This distributive aspect of sensory processing not only grants evolutionary advantages, such as prompt reactions to threats, but also serves to reduce the uncertainty derived from individual sensory estimates and to endure brain damage or sensory loss (van Atteveldt et al., 2014).

Previous research on audio-visual integration in humans has primarily relied on task-based paradigms employing simple stimuli (e.g., checkerboards or flashes as visual stimuli, and tones or noises as auditory information) [for a review see, (Murray et al., 2016)], limiting the generalizability of findings to complex, real-life situations (Hari et al., 2015; Khosla et al., 2021; Varoquaux & Poldrack, 2019). Functional magnetic resonance imaging (fMRI) experiments during naturalistic film viewing provide an alternative approach, enabling the study of audio-visual integration in ecologically valid contexts and facilitating investigations into high-level processes, such as social cognition (Finn, 2021).

During film viewing, we effortlessly recognise and follow the mental states of onscreen characters, a process associated with theory-of-mind function (Abell et al., 2000; Castelli et al., 2000; Saxe & Wexler, 2005). This seemingly undemanding task, however, can be complex; especially when visual cues are ambiguous. In these cases, viewers strongly rely on contextual cues, such as the accompanying music, to aid their comprehension (Boltz, 2001; Cohen, 2005, 2013). When integrated into a visual context such as film, music’s influence goes beyond mirroring a meaning portrayed by the visual images (Chion, 1994; Pudovkin, 1929). Music can shape our experience of the film’s narrative, affecting perceptual judgements, emotion and memory of the events (Bolivar et al., 1994; Boltz, 2001; Boltz et al., 1991; Bullerjahn & Güldenring, 1994; Cohen, 2001, 2005, 2013; Hoeckner et al., 2011; Sirius & Clarke, 1994; Tan et al., 2007).

Despite the significant impact of music on film, there is limited neuroscientific research investigating the joint processing of music and visual information. Previous studies have shown that the combined presentation of music and visual information increases activation in brain structures associated to emotion (Baumgartner et al., 2006; Eldar et al., 2007). However, the lack of control over specific musical structure variables in these studies has made it challenging to determine how such features modulate brain function.

In the present study, we employed fMRI during naturalistic film viewing to examine the neural mechanisms underlying our emotional responses to musical dissonance (Bigand et al., 1996; Bigand & Parncutt, 1999; Huron, 2008; Lerdahl & Krumhansl, 2007). Although one of the most common harmonic manipulations in film music (e.g., Spielberg’s *Jaws* or Kubrick’s *The Shining*), systematically controlled transformations of tonal dissonance have not yet been neuroscientifically studied in regard to affective processing biases of visual contexts. By strictly controlling musical variables such as instrumental timbre, dynamics, rhythm, textural density and melodic contour, we aimed to investigate the effects of tonal dissonance on the neural processing of emotional valence during mental state inferences (Van Overwalle, 2009; Van Overwalle et al., 2009). Specifically, we focused on valence judgments during mental state attribution to the main character in an animated short film (see methods, Figure 7).

The assumption was that although participants would watch the same visual scene, tonal dissonance would bias their interpretative framework (Boltz, 2001) leading to more negatively-valenced mental state inferences. Furthermore, we predicted that early visual processing would interact with higher-level auditory association areas, indicating the influence of music on the visual experience. Accordingly, we hypothesised that the auditory ventral stream (Belin et al., 2000; Hickok & Poeppel, 2000; Kaas & Hackett, 1999), a brain pathway involved in mapping contextual sound cues to semantic associated attributes, would modulate low-level visual encoding via feedback projections.

To test our hypotheses, we designed an experimental paradigm that instructed participants to ascribe mental states to the main character in the film clip. Through investigating the recruitment of theory-of-mind neural substrates, the influence of tonal dissonance on mental state inferences, the interaction between early visual processes and sound-to-meaning systems, and the potential impact of top-down contextual influences on visual encoding in V1, we aimed to uncover the cognitive mechanisms and the neural underpinnings of our emotional responses to musical dissonance during naturalistic film viewing.

## Results

### Behavioural

#### Impact of tonal consonance/dissonance on valence judgements

During the present fMRI study, thirty-eight subjects watched the same short-film with controlled consonant or dissonant music, in randomized order. Prior to the start of the audio-visual film clip subjects were instructed to “*think about the intentions of the main character in the following film clip*”. Following the clip participants were asked to rate the valence of the movie character’s intentions on a scale ranging from one (good intentions) to four (bad intentions). Two control conditions were included; i) visual alone category (i.e., film clip with no soundtrack but same instruction as above) and, ii) film clip with an instruction to describe the “physical” appearance of the character, to control for multimodal sensory processing, working memory and attentional demands of the task, without cueing subjects to attend specifically to mental states (see methods and Figure 7 for details).

Behavioural results revealed significant differences between the consonant, dissonant and visual alone conditions (Table 1). A repeated measures MANOVA showed a significant effect of condition on the judgements of valence regarding the character’s intentions (*Wilks’ Lambda* = *0.509, F* (_2, 36_) = 17.348, *p* < 0.001). [The data did not violate the sphericity assumption; Mauchly’s test of sphericity was not significant (*p* = 0.345)]. Post hoc tests (Bonferroni corrected) indicated that the average valence rating for the dissonant condition was significantly more negative than the valence rating for the consonant condition (*p* < 0.001, *d* = 0.895) and the visual alone condition (*p* = 0.013, *d* = 0.368). There was also a significant difference in valence ratings between the consonant and the visual alone conditions (*p* < 0.001, *d* = 0.526). Concretely, subjects ascribed significantly more negative (positive) intentions to the character on-screen when the film clip was paired with dissonant (consonant) music. Polynomial contrasts on the mean ratings for the three categories indicated a significant linear trend (*F* (_1, 5.263_) = 15.289, *p* < 0.001), dissonance > visual alone > consonance (Table 1).

**Table 1.**
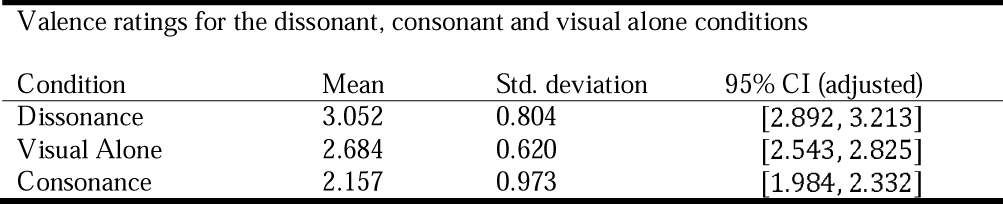
Valence means with standard deviations and 95% confidence intervals for each condition (consonance: movie with consonant music; dissonance: movie with dissonant music, visual alone: movie without music). The range for valence ratings was one (good intentions) to four (bad intentions).

| Valence ratings for the dissonant, consonant and visual alone conditions |  |  |  |
| --- | --- | --- | --- |
| Condition | Mean | Std. deviation | 95% CI (adjusted) |
| Dissonance | 3.052 | 0.804 | [2.892, 3.213] |
| Visual Alone | 2.684 | 0.620 | [2.543, 2.825] |
| Consonance | 2.157 | 0.973 | [1.984, 2.332] |

The results support previous research that has examined the affective reactions induced by consonance/dissonance (Blood et al., 1999; Costa et al., 2000; Fritz et al., 2009, 2013; Koelsch et al., 2006; Plomp & Levelt, 1965; Trainor & Heinmiller, 1998); and, specifically converge with studies assessing the impact of consonance/dissonance on valence judgements during mental state attribution, in which increasing levels of dissonance induce more negatively-valenced inferences (Bravo, 2013; Bravo, Cross, Hawkins, et al., 2017; Bravo, Cross, Stamatakis, et al., 2017).

### Functional MRI

#### BOLD signal changes – subtractive analysis

##### Effects of the theory-of-mind task compared with the control condition (physical appearance)

The comparison between the audio-visual theory-of-mind (ToM) task *(i.e., “think about the intentions of the main character”)* against the audio-visual control condition (“*focus on the physical appearance of the main character”*) evidenced signal changes in bilateral supramarginal and inferior parietal gyrus (Table 2A, Figure 1d). The results are in line with previous studies that have assessed the neural systems supporting implicit non-verbal ToM encoding (Grosse Wiesmann et al., 2020) and corroborate that our task engaged mental state inference processing (Van Overwalle, 2009; Van Overwalle et al., 2009).

**Fig. 1.**
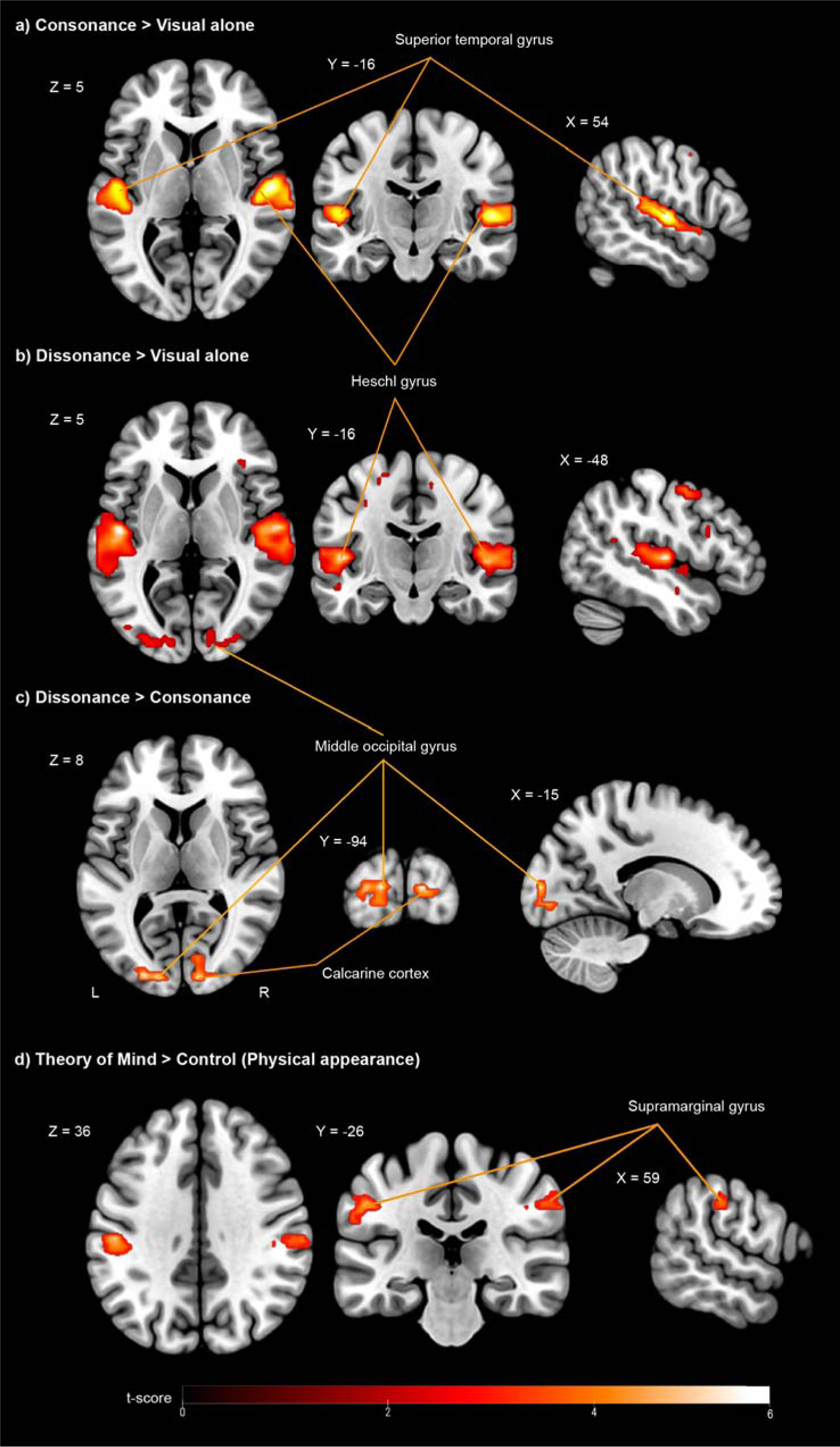
Effects of tonal dissonance and theory-of-mind function during naturalistic film-viewing. fMRI results (FWE-corrected *p* < 0.05 for cluster-level inference). Coloured areas (red) reflect statistical parametric maps (SPM) superimposed onto a standard brain in stereotactic MNI space. a) consonance > visual alone and b) dissonance > visual alone: similar patterns of brain response were observed bilaterally in primary and secondary auditory cortices including Heschl’s gyri. The contrast dissonance > visual alone further evidenced signal changes in occipital areas. c) Statistical parametric maps showing voxels in the middle occipital gyrus and calcarine cortex in which the response was higher during the dissonant compared to consonant condition. d) Statistical parametric maps showing activation in supramarginal and inferior parietal gyrus (bilaterally) for the comparison between ToM > control conditions.

**Table 2.** (A) fMRI Results (at threshold *p* < 0.001 voxel-level uncorrected, *p* < 0.05 cluster-level FWE-corrected) of group General Linear Model for the contrasts: dissonance > consonance, consonance > dissonance, consonance > visual alone, dissonance > visual alone, and audiovisual task (Theory of Mind) > audiovisual control (Physical appearance). (B) Results of group psychophysiological interactions analysis (PPI) with seed voxels (spheres with a 6 mm radius) located around highest peak for each subject within the right calcarine cortex and within the left middle occipital gyrus (contrast dissonance > consonance). The regions described showed stronger positive functional connectivity with either of these regions during the dissonant condition compared to the consonant condition. (C) Dynamic Causal Modeling (DCM) Bayesian model selection (BMS) results: Conditional probability (expected posterior probability representing the probability of a model given the observed data) and exceedance probability (probability compared with other tested models). Model 1 specified a connection from the rMTG to the lMOG (hypothesized). Model 2 specified a connection from the lMOG to the rMTG. Model 3 specified bidirectional connections between both regions. Model 1 obtained the most evidence. Abbreviations: L: left; R: right; TPJ: Temporo-parietal junction: supramarginal gyrus, angular gyrus and superior temporal gyrus; mPFC: medial prefrontal cortex, ACC, anterior cingulate cortex Cl.: areas integrating the above detailed cluster-level p-value.

| Region | Cluster Peak MNI | Voxels | Max t-value (z-value) | Mean t (std.) | p-value (FWE) |
| --- | --- | --- | --- | --- | --- |
| **(A) Subtractive analysis** |  |  |  |  |  |
| *Dissonance > Consonance* |  |  |  |  |  |
| Occipital Mid L | -15 -94 8 | 78 | 5.70 (4.80) | 4.05 (0.47) | < 0.0001 |
| Occipital Sup L |  | 17 |  | 4.07 (0.64) | Cl. |
| Occipital Inf L |  | 15 |  | 4.23 (0.58) | Cl. |
| Lingual L |  | 8 |  | 3.82 (0.49) | Cl. |
| Calcarine L |  | 5 |  | 3.85 (0.19) | Cl. |
| Calcarine R | 12 -94 8 | 37 | 5.51 (4.80) | 3.72 (0.33) | < 0.037 |
| Cuneus R |  | 5 |  | 4.72 (0.81) | Cl. |
| Occipital Sup R |  | 5 |  | 3.79 (0.45) | Cl. |
| Occipital Mid R |  | 4 |  | 3.60 (0.16) | Cl. |
| *Consonance > Dissonance* |  |  |  |  |  |
| (No suprathreshold activations were observed) |  |  |  |  |  |
| *Consonance > Visual Alone* |  |  |  |  |  |
| Temporal Sup R | 54 -16 5 | 237 | 8.31 (6.21) | 4.76 (1.05) | < 0.0001 |
| Heschl R |  | 44 |  | 4.69 (1.18) | cl. |
| Temporal Sup L | -48 -25 8 | 177 | 7.05 (5.58) | 4.58 (0.87) | < 0.0001 |
| Heschl L |  | 25 |  | 5.02 (1.09) | cl. |
| *Dissonance > Visual Alone* |  |  |  |  |  |
| Temporal Sup L | -48 -16 5 | 296 | 8.54 (6.31) | 4.74 (0.99) | < 0.0001 |
| Heschl L |  | 42 |  | 5.29 (1.34) | cl. |
| Temporal Sup R | 60 -13 5 | 399 | 8.14 (6.12) | 4.78 (1.00) | < 0.0001 |
| Heschl R |  | 56 |  | 5.19 (1.01) | cl. |
| Occipital Inf L | -33 -88 -10 | 96 | 6.76 (5.42) | 4.34 (0.74) | < 0.0001 |
| Occipital Mid L |  | 86 |  | 4.33 (0.81) | cl. |
| Occipital Sup L |  | 10 |  | 3.78 (0.28) | cl. |
| Calcarine L |  | 3 |  | 4.53 (0.20) | cl. |
| Precentral L | -39 -7 62 | 144 | 6.63 (5.35) | 4.20 (0.74) | < 0.0001 |
| Supp Motor Area L |  | 84 |  | 3.93 (0.51) | cl. |
| Occipital Inf R | 45 -73 -7 | 38 | 5.47 (4.65) | 3.81 (0.37) | < 0.0001 |
| Calcarine R |  | 37 |  | 3.68 (0.29) | cl. |
| Occipital Mid R |  | 20 |  | 3.58 (0.20) | cl. |
| Occipital Sup R |  | 2 |  | 3.72 (0.06) | cl. |
| Parietal Sup L | -30 -52 53 | 69 | 5.32 (4.56) | 3.79 (0.37) | < 0.0001 |
| Parietal Inf L |  | 30 |  | 4.13 (0.46) | cl. |
| Frontal Inf Tri L | -42 17 26 | 34 | 5.01 (4.38) | 3.94 (0.44) | 0.006 |
| Frontal Inf Oper L |  | 10 |  | 3.53 (0.18) | cl. |
| Insula L |  | 5 |  | 3.76 (0.37) | cl. |
| Frontal Inf Tri R | 54 32 11 | 52 | 4.82 (4.22) | 3.74 (0.36) | 0.008 |
| Frontal Inf Oper R |  | 6 |  | 3.95 (0.47) | cl. |
| Insula R |  | 1 |  | 3.35 (-) | cl. |
| Mid Prefrontal R | 27 41 20 | 17 | 4.79 (4.20) | 3.75 (0.43) | < 0.0001 |
| Cingulum Ant R |  | 13 |  | 3.88 (0.34) | cl. |
| *Task (Theory of Mind) > Control (Physical appearance) - ROI analysis: TPJ, mPFC, ACC* |  |  |  |  |  |
| Parietal Inf L | -57 -28 29 | 100 | 6.36 (5.20) | 4.17 (0.58) | < 0.0001 |
| SupraMarginal L |  | 43 |  | 4.16 (0.72) | cl. |
| Postcentral L |  | 19 |  | 3.92 (0.40) | cl. |
| Parietal_Inf_R | 33 -37 41 | 25 | 5.44 (4.63) | 3.90 (0.38) | < 0.0001 |
| SupraMarginal_R |  | 10 |  | 4.39 (0.55) | cl. |
| Postcentral R |  | 9 |  | 4.02 (0.29) | cl. |
| **(B) Functional connectivity analysis (PPI)** |  |  |  |  |  |
| Seed region (6 mm radius sphere) located around highest peak for each subject within the **right Calcarine** |  |  |  |  |  |
| *PPI – dissonance > consonance* |  |  |  |  |  |
| Temporal Mid R | 51 -70 -4 | 182 | 7.86 (5.99) | 4.45 (0.92) | < 0.0001 |
| Temporal Inf R |  | 81 |  | 4.76 (1.09) | cl. |
| Occipital Mid R |  | 27 |  | 4.39 (0.83) | cl. |
| Occipital Mid L | -42 -76 -1 | 94 | 5.40 (4.61) | 4.16 (0.56) | < 0.0001 |
| Temporal Mid L |  | 60 |  | 4.05 (0.55) | cl. |
| Temporal Inf L |  | 3 |  | 3.59 (0.21) | cl. |
| Seed region (6 mm radius sphere) located around highest peak for each subject within the **left middle Occipital** |  |  |  |  |  |
| *PPI – dissonance > consonance* |  |  |  |  |  |
| Temporal Mid R | 51 -70 -1 | 88 | 6.33 (5.18) | 4.29 (0.82) | < 0.0001 |
| Temporal Inf R |  | 27 |  | 4.01 (0.54) | cl. |
| Occipital Mid R |  | 9 |  | 4.15 (0.50) | cl. |
| Occipital Mid L | -48 -76 5 | 62 | 5.77 (4.84) | 4.18 (0.64) | < 0.0001 |
| Temporal Mid L |  | 60 |  | 3.96 (0.55) | cl. |
| Temporal Inf L |  | 3 |  | 3.79 (0.17) | cl. |
| **(C) Effective connectivity analysis (DCM)** |  |  |  |  |  |
| DCM – BMS (Effects of dissonance) Models | | | | Expected sk\|Y | Exceedance $\psi_k$ |
| 1) right middle temporal gyrus → left middle occipital gyrus |  |  |  | 0.6874 | 0.9104 |
| 2) left middle occipital gyrus → right middle temporal gyrus |  |  |  | 0.1499 | 0.0350 |
| 3) Bidirectional |  |  |  | 0.1627 | 0.0546 |

##### Effects of music (in an audio-visual context) compared to visual alone

To establish the general effects of music compared to visual (alone) information, we contrasted consonance > visual alone and dissonance > visual alone. These comparisons showed similar patterns of brain response in primary and secondary auditory cortices (AC: superior temporal gyrus, Heschl’s gyrus) bilaterally (Table 2A, Figure 1a,b). These findings converge with previous evidence from studies that have employed music listening paradigms containing harmonized chord progressions (Brown et al., 2004; Menon, 2002; Ohnishi et al., 2001), in which signal changes were observed bilaterally in primary and secondary AC when comparing sound conditions against no-sound (or silent) conditions. The contrast between dissonance > visual alone unveiled additional activations in the visual cortex (inferior, middle, superior occipital gyrus and bilateral calcarine cortices), which we interpret as effects driven by increased tonal dissonance, and not by sound information per se (see following subsection).

##### Effects of musical dissonance compared to musical consonance within an audiovisual context

The contrast of interest, dissonance > consonance, revealed significant signal changes in the visual cortex (bilateral middle and superior occipital gyrus, left inferior occipital gyrus), including the primary visual cortex (bilateral calcarine cortex) (Table 2A, Figure 1c). The opposite contrast (consonance > dissonance) did not show any suprathreshold signal changes. Given the experimental design, showing identical visual information in both conditions, the results substantiate our prediction of early visual systems modulated by contextual sound cues and indicate a role of the visual system beyond visual recognition (Bullier, 2001; Gilbert & Li, 2013; Petro et al., 2017). To our knowledge this is the first report of activations encompassing early visual cortices (V1) modulated by musical information with systematically controlled manipulations of tonal consonance/dissonance level.

#### Functional connectivity

##### Psychophysiological interaction (PPI) analysis

The function of the primary visual cortex in visual recognition is known to be strongly modulated by multisensory information (De Meo et al., 2015; Ghazanfar et al., 2005; Murray et al., 2016; van Atteveldt et al., 2014). In particular, auditory feedback signals are the largest contributor (Murray et al., 2016; Shams et al., 2000). To further elucidate why dissonance recruited additional regions and to examine potential neuromodulatory influences on early visual processing, we measured functional connectivity with PPI (Friston et al., 1997). A PPI analysis identifies voxels that increase their relationship with a seed region of interest (ROI) in a given psychological context, such as when participants watch the film with dissonant compared to consonant music. Seeds ROIs (spheres with a 6 mm radius) were defined around the highest activated peak for each subject in the left middle occipital gyrus and in the right calcarine cortex, based on the significantly activated clusters for the *contrast dissonance > consonance*. The results (displayed in Table 2B, Figure 2) showed that, during the dissonant condition (compared with the consonant condition), a cluster with peak in the right middle (extending to the right inferior) posterior temporal gyrus exhibited stronger functional connectivity with the right calcarine cortex and with the left middle occipital gyrus, indicating that tonal dissonance modulated the connection between early visual encoding and the ventral auditory system [represented by the right posterior middle temporal gyrus; (Belin et al., 2000; Kaas & Hackett, 1999; Rauschecker, 1998; Romanski et al., 1999)], which is known to serve as a sound-to-meaning interface, mapping sound-based cues (e.g., distinct levels of dissonance) to their semantic associated attributes (e.g., emotional valence).

**Fig. 2.**
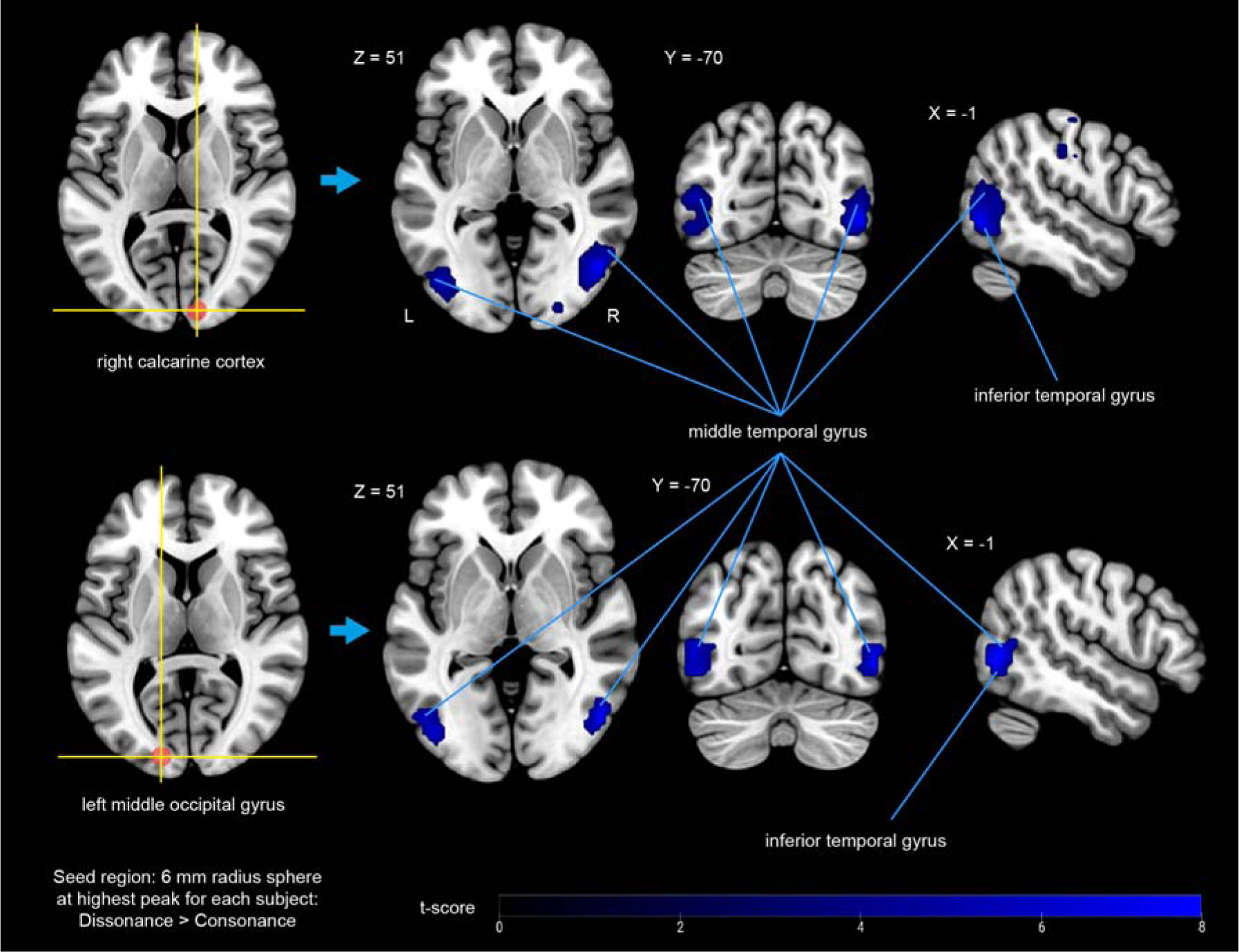
Functional connectivity modulated by tonal dissonance. fMRI results (FWE-corrected *p* < 0.05 for cluster-level inference) Psycho-physiological interaction analysis (PPI). Blue colour identifies voxels in the middle (extending to inferior) posterior temporal gyrus, which exhibited stronger functional connectivity with the primary visual cortex (PPI seed regions were 6mm spheres around the highest activated peak for each subject within the right calcarine cortex and the left middle occipital gyrus, for the contrast dissonance > consonance).

##### Dynamic Causal Modeling (DCM) analysis (effective connectivity)

To further examine the causal flow of information between visual processing and the ventral auditory system and, specifically, to assess whether dissonance engages top-down or bottom-up interactions, we carried out an effective connectivity analysis using dynamic causal modelling (DCM) (Friston, 2003). We hypothesized that distinct dissonance/consonance levels would exert differential influences on early visual processing through feedback projections from the ventral auditory stream (Hickok & Poeppel, 2007). To test this hypothesis, DCM analysis was conducted for each subject on two ROIs: i) the left middle occipital gyrus (lMOG), which showed the strongest activation in the subtractive analysis *dissonance > consonance;* and *ii)* the right middle temporal gyrus (rMTG), which exhibited the most robust functional coupling with the lMOG (see methods for detailed information). The modulatory effect of interest was the dissonance condition, since this category appeared to strongly modulate the functional interaction (PPI) between the aforementioned areas. We modelled the direction of interaction. For each subject, three models were defined. Model 1 specified a connection from the rMTG to the lMOG (hypothesized). Model 2 specified a connection from the lMOG to the rMTG. Model 3 specified bidirectional connections between both regions. Model 1, in which the left middle occipital gyrus received information from the right middle temporal gyrus obtained the most evidence. Bayesian model averaging further indicated a negative modulation effect of dissonance (−0.0002) on the connection from the rMTG to the lMOG that changed the strength of the intrinsic connection (0.0189) (Table 2C). The results are consistent with a role of feedback signals being sent from the auditory ventral stream to the early visual cortex, modulated by increased tonal dissonance.

## Discussion

### Impact of tonal consonance/dissonance on valence judgements during social cognition processing

The present study aimed to investigate the cognitive and neural mechanisms underlying music’s effect on valence inferences during mental state attribution. To assess the capacity of our task to recruit mentalizing neural substrates we compared the audio-visual *theory-of-mind* condition with the audio-visual *control* condition (focused on physical appearance). This comparison revealed signal changes in bilateral supramarginal and inferior parietal gyrus; areas supporting implicit non-verbal ToM reasoning (Grosse Wiesmann et al., 2020) as well as emotional and visual perspective taking, action observation, social attention and encoding biases (Kanske et al., 2016; Schurz et al., 2015; Silani et al., 2013; Steinbeis et al., 2015).

Following each film sequence, participants were asked to rate the valence of the movie character’s intentions (i.e., positive or negative). The results showed that musical dissonance led to significantly more negative mental state inferences, exposing its strong emotional impact. Whilst in line with previous research that has examined the negative affects elicited by dissonance (Blood et al., 1999; Costa et al., 2000; Fritz et al., 2009, 2013; Koelsch et al., 2006; Plomp & Levelt, 1965; Trainor & Heinmiller, 1998), our findings extend the evidence for its strong influence on valence judgements during mental state attribution (Bravo, 2013; Bravo, Cross, Hawkins, et al., 2017; Bravo, Cross, Stamatakis, et al., 2017).

### Activation of V1 by auditory cues with increased level of tonal dissonance

Univariate whole-brain analysis revealed significant bilateral activation of primary visual cortices (calcarine cortex) including the middle and superior occipital gyrus, while participants were watching the film sequence accompanied by dissonant, compared to consonant, music. The results endorse our prediction of modulations in early visual systems driven by musical information and provide, to our knowledge, the first demonstration for the engagement of crossmodal low-level sensory substrates in response to tonal dissonance.

The findings are in agreement with previous models which have highlighted the adaptable properties of cortical neurons (Bullier, 2001; Gilbert & Li, 2013; Petro et al., 2017). Electrophysiological recordings show that V1 neurons function as spatio-temporal filters encoding elementary visual features, upon which feedforward connections with higher visual areas assist in representing progressively more complex aspects of the visual scenario (DiCarlo et al., 2012). Although the primary function of V1 is visual perception, its role expands beyond visual recognition and encompasses multisensory aspects (De Meo et al., 2015; Ghazanfar et al., 2005; Murray et al., 2016; van Atteveldt et al., 2014), with the largest contribution of crossmodal re-entrant signals being auditory feedback (Murray et al., 2016). An illustration at relatively basic sensory processing levels has been reported by Shams et al. (2000). The authors showed how a single visual flash could be misperceived as multiple flashes when paired with multiple auditory beeps. Our results go beyond this previous work by demonstrating that dissonant sound cues can qualitatively bias higher-level mental state inference processes, by exerting neuromodulatory effects on early visual encoding. The findings emphasise the role of the brain as a highly distributed processing engine in which areas across multiple hierarchies work in parallel to perform complex cognitive functions.

### Top-down modulation of visual processing

We investigated whether V1 could be subject to top-down contextual influences modulated by tonal dissonance. Psychophysiological interaction analysis showed strong coupling between the right middle posterior temporal gyrus and V1 in response to dissonance. Effective connectivity analysis demonstrated that musical dissonance modulated visual processing via top-down feedback inputs from the right middle posterior temporal gyrus to V1.

The prevailing view of sensory information processing is based on feedforward connections. Within the visual system these type of connections characterise a hierarchy of cortical areas initiated in V1 and ascending via the ventral or dorsal pathways (Mishkin et al., 1983; Ungerleider, 1994). Each feedforward connection, however, coexists with a reciprocal feedback connection that carries contextual information. Cortical neurons have been described as ‘active blackboards’ or ‘adaptive processors’ (Bullier, 2001; Gilbert & Li, 2013), which can modify their function and response according to the behavioural context or the specific demands of the task being carried out. It is well established that the visual cortex can be subjected to diverse top-down influences, including among others, those of attention, reward, and emotion responses (Petro et al., 2014). Top-down signals represent influences from higher-order cognitive representations that impact earlier stages of information processing. These higher-order representations can carry contextual information that prepares the visual system for optimizing behavioural responses (Desimone & Duncan, 1995; Gilbert & Li, 2013).

While prior research has examined the functional effects of auditory information in the primary visual cortex, most of these studies have employed basic stimuli or simplified displays to investigate the role of sound in assisting the spatial localization of visual inputs, consequently reducing the complexity of naturalistic interaction conditions (Peelen & Kastner, 2014). Only few studies have assessed higher-order cognitive representations using naturalistic stimulus. Petro et al. (2013) employed auditory information paired with uniform visual stimulation (i.e., a blank fixation screen). Using multivoxel pattern analysis (MVPA), the authors showed that sound imagery could be decoded from the early visual cortex. However, to date, there is still no evidence of auditory information readouts from V1 when it receives a more driving visual stimuli (Petro et al., 2017). We hereby broaden these previous findings by using non-uniform, conspicuous visual stimuli (i.e., naturalistic film-viewing), and demonstrating a strong response of the primary visual cortex to musical dissonance using univariate whole-brain analyses.

Linking the schema theory with predictive coding approaches (Bar, 2009; Friston, 2005, 2009), we propose a potential mechanism to explain these modulatory influences. We argue that our manipulation of the musical stimuli triggered high-level contextual representations, which were fed back to early visual cortices to facilitate the interpretation of the visual information. Throughout an audio-visual experience such as watching a film, musical affect can direct the viewer’s attention toward visual aspects that portray a matching connotative meaning. This, in turn, can allow the spectator to develop certain inferences about the characters’ internal motivations, e.g., their intentions, therefore contributing to the comprehension of the story. At a behavioural level, this mechanism has been conceptualized as “mood congruency effects” and explained by means of the schema theory (Boltz, 2001). Schemas (Mandler & Johnson, 1976) [also described as scripts (Schank, 1975) or frames (Minsky, 1974)] are memory associations built by extracting repeating patterns and statistical regularities from our environment. This “related” information refers to events that tend to be linked on some level (such as dissonance leading to negative valence), and is thought to be clustered in memory structures that serve as the building block for predictions (Bar, 2004, 2009; Bar & Ullman, 1996). From a socio-cognitive perspective, schemas allow us comprehend the social environment: they help us understand why people behave in the ways they do, and predict what is most likely to happen next. We posit that the predictions set by these memory associations are probably manifest in the signal changes we observe in the primary visual cortex.

### Feedback influences from the auditory “what” ventral stream (mapping sound cues to meaning) to V1

The two-streams hypothesis is a model of the neural processing of the cortical organization of vision (Goodale & Milner, 1992). The model proposes two information processing pathways originating in the occipital cortex. A dorsal visual stream leading to the parietal lobe, which is involved in processing spatial relationships among objects ("where pathway"); and a ventral visual stream that projects to the inferior temporal cortex and which is crucial for the visual identification of objects (also known as the "what pathway").

Intersecting Goodale and Milner’s model with Wernicke’s research (Wernicke, 1874), which suggested that sensory representations of speech should at least interface with two systems: a motor-articulatory and a conceptual system; Hickok and Poeppel (2004) advanced an analogous dual stream hypothesis for auditory language processing. The model entails a ventral stream supporting conceptual representations via projections to the middle posterior temporal gyrus, and a dorsal stream entailing motor representations via projections to temporal-parietal regions (Hickok & Poeppel, 2000, 2007). Although the specific role of the dorsal auditory stream is still a subject of debate with various hypotheses including localization, spectral processing and auditory-motor integration, there is general agreement on the *“what”* processing function of the auditory ventral stream (Belin et al., 2000; Hickok & Poeppel, 2000; Kaas & Hackett, 1999) For instance, the prototypical verbal paradigm for targeting the ventral auditory pathway involves listening to meaningful speech versus meaningless pseudo speech (Saur et al., 2008). The ventral pathway serves as a sound-to-meaning interface by mapping sound-based representations to distributed conceptual representations (Belin et al., 2000; Kaas & Hackett, 1999; Rauschecker, 1998; Romanski et al., 1999). In our study, functional/effective connectivity analysis revealed a causal flow of information from the right middle posterior temporal gyrus to V1 modulated by tonal dissonance. These findings suggest the involvement of the auditory ventral stream in encoding of the above-described schemas: building global representations for perceptual and semantic associated attributes by mapping non-verbal contextual sound cues to meaning (e.g., dissonant music leading to negative valence). These, in turn, provide an internal model or interpretative framework that helps to organize the visual experience into a comprehensible whole via feedback projections to the early visual cortices (Figure 3).

**Fig. 3.**
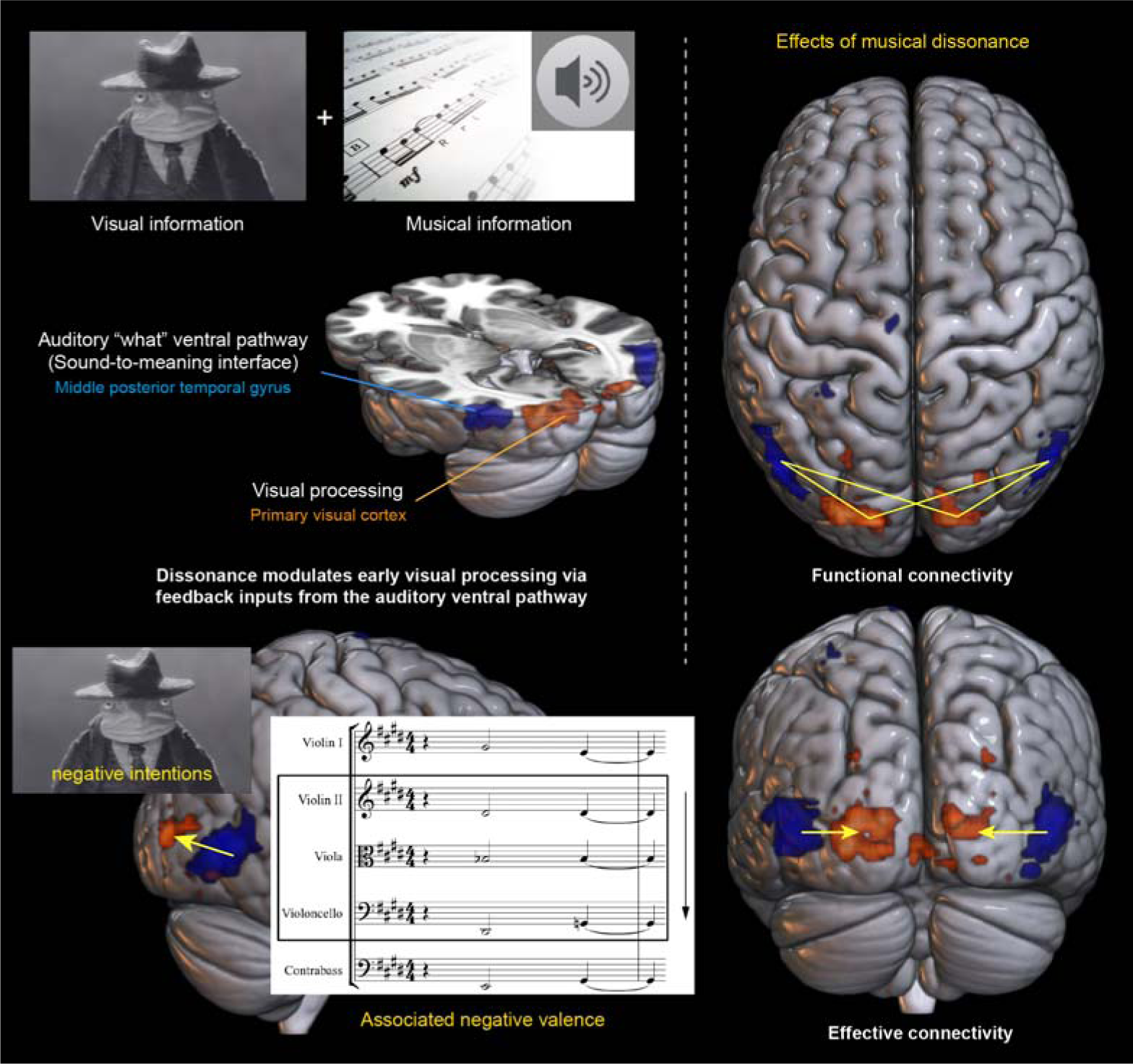
Schematic representation of the findings. Left panel: The auditory “what” ventral stream plays a role in assigning meaning to non-verbal sound cues, such as dissonant music conveying negative emotions, providing an interpretative framework that serves to process the audio-visual experience. Right panel: Functional/effective connectivity analysis showed a coupling between the auditory ventral stream (AVS) and V1 in response to tonal dissonance, and demonstrated the modulation of early visual processing via top-down feedback inputs from the AVS to V1.

Why does tonal dissonance, rather than consonance, drive stronger the modulatory effects on V1?

It has been consistently acknowledged that emotionally laden stimuli benefit from preferential processing given their adaptive significance (Carretié et al., 2001; Dolcos et al., 2004, 2020; Dolcos & Cabeza, 2002; Kensinger & Schacter, 2008; Kosslyn et al., 1996; Lang et al., 1998; Mickley Steinmetz & Kensinger, 2009; Pourtois et al., 2013; Vuilleumier, 2015; Vuilleumier et al., 2004). We have previously shown that the specific harmonic manipulation applied in this study can isolate emotional valence from the arousal and potency dimensions, facilitating a novel access to the neural representation of negative emotion (Bravo, 2013; Bravo et al., 2019). Negative valence can further modulate the interaction between visual processing and attentional systems implicated in the appraisal of behaviourally relevant, unexpected and potentially threatening events (Bravo, Cross, Hawkins, et al., 2017; Bravo et al., 2019; Corbetta et al., 2008; Corbetta & Shulman, 2002). We postulate that the negative valence associated with the dissonant musical background can explain the heightened weight on visual sensory evidence. Our findings conceptually expand previous work, which has exclusively focused on the affective value associated with visual stimuli, to provide evidence of crossmodal interactions and enhanced neural responses in early visual pathways signalled by non-verbal auditory information: music.

The ultimate purpose of our work is to develop advanced methods of applying musical information in naturalistic paradigms to characterize complex mental health disorders. Relative to traditional paradigms, naturalistic stimuli, such as the one used in this study, are often more engaging than conventional highly-constrained tasks, and therefore allow the observation of brain activity that closely resembles to freeform cognition. Film-watching provides an effective means to study multiplexed neural responses, from low-level sensory processes to high-level components: we showed that a simple, but controlled, manipulation of musical dissonance can elucidate the brain’s sound-to-meaning interface and its distributive effects during social cognition. We envision the use of non-verbal music-based fMRI paradigms for identifying behavioural signatures and measurable neural markers associated with severe psychopathology.

## Conclusions

In the present study, we employed fMRI to assess the effects of musical dissonance upon the affective processing of visual information during naturalistic film viewing. Participants watched the same short-film with either dissonant or consonant music. Compared to consonant music, dissonance led to more negative mental state attributions. Neuroscientific analyses revealed that tonal dissonance modulated early visual processing via top-down cortical feedback inputs from the auditory “what” ventral stream to the primary visual cortex. We offer evidence for the involvement of the auditory ventral stream in mapping non-verbal sound cues to meaning, providing an internal model that serves to organize the audio-visual experience. Taken together, these findings demonstrate the critical role of multisensory processing, particularly audio-visual integration, in shaping higher-order functions such as social cognition.

## Materials and Methods

### Subjects

Data for the fMRI experiment were obtained from thirty-eight healthy volunteers. The participants (18 females and 20 males; mean age = 27±4) were native German speakers, with no history of neurological or psychiatric illness, or use of psychotropic medication. They reported no long-term hearing impairment. All participants were right-handed. The sample did not include any professional musician. Five participants (2 male, 3 female) reported having received informal musical training for less than three years. All subjects gave written informed consent. The study received ethical approval from the Ethics Commission at TU-Dresden (Reference Number: EK 305072016).

### Stimulus Material

The task was constructed with the purpose of assessing affective mental state inferences (Happé, 2003; Schultz et al., 2004; Van Overwalle, 2009) biased by musical structural features (Bravo, Cross, Hawkins, et al., 2017), and to enable the investigation of the underlying neural encoding. The paradigm was built in the form of an audio-visual film clip, based on the short film “Man with pendulous arms” (1997, directed by Laurent Gorgiard) (Figure 4).

**Fig. 4.**
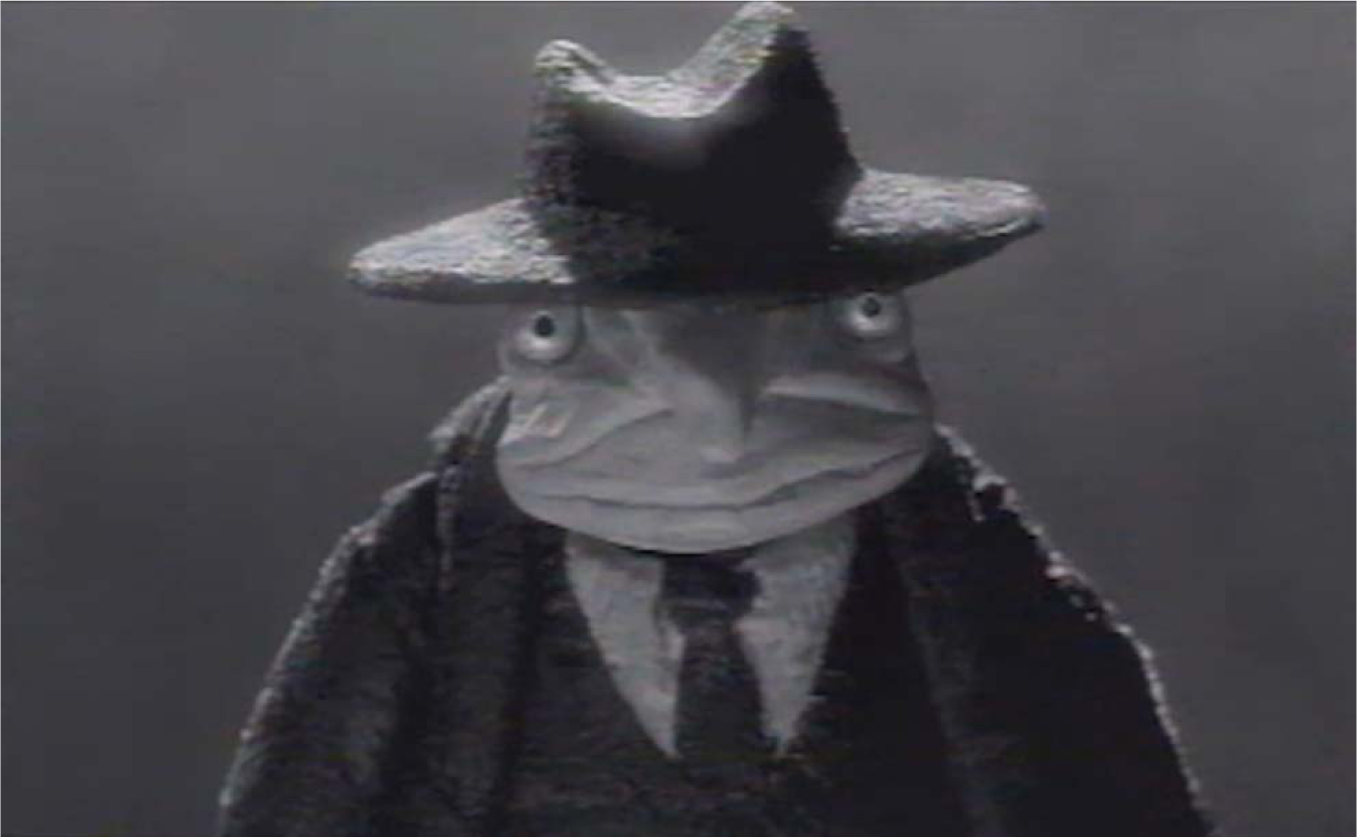
Screenshot from the short film “Man with pendulous arms” (1997, Laurent Gorgiard) employed in the fMRI experiment. Participants were instructed to “*think about the intentions of the main character*”.

The materials employed in the present study were created for, and tested in two a previous experiments (Bravo, 2013; Bravo et al., 2019). The musical stimuli comprised two soundtracks (i.e., consonant and dissonant) composed for the same short film. In summary, a choral piece, written in the form of musical variations, was made to sound more or less consonant or dissonant by modifying its harmonic structure (i.e., interval content manipulation) producing two otherwise-identical versions of the same musical piece. The consonant condition consisted of a theme followed by three variations written by FB in a romantic musical style. The dissonant condition was achieved by lowering by a semitone the second violin, viola and violoncello lines (of the consonant piece), while maintaining the other instruments (i.e., first violin and double bass) at their original pitch (Figure 5).

**Fig. 5.**
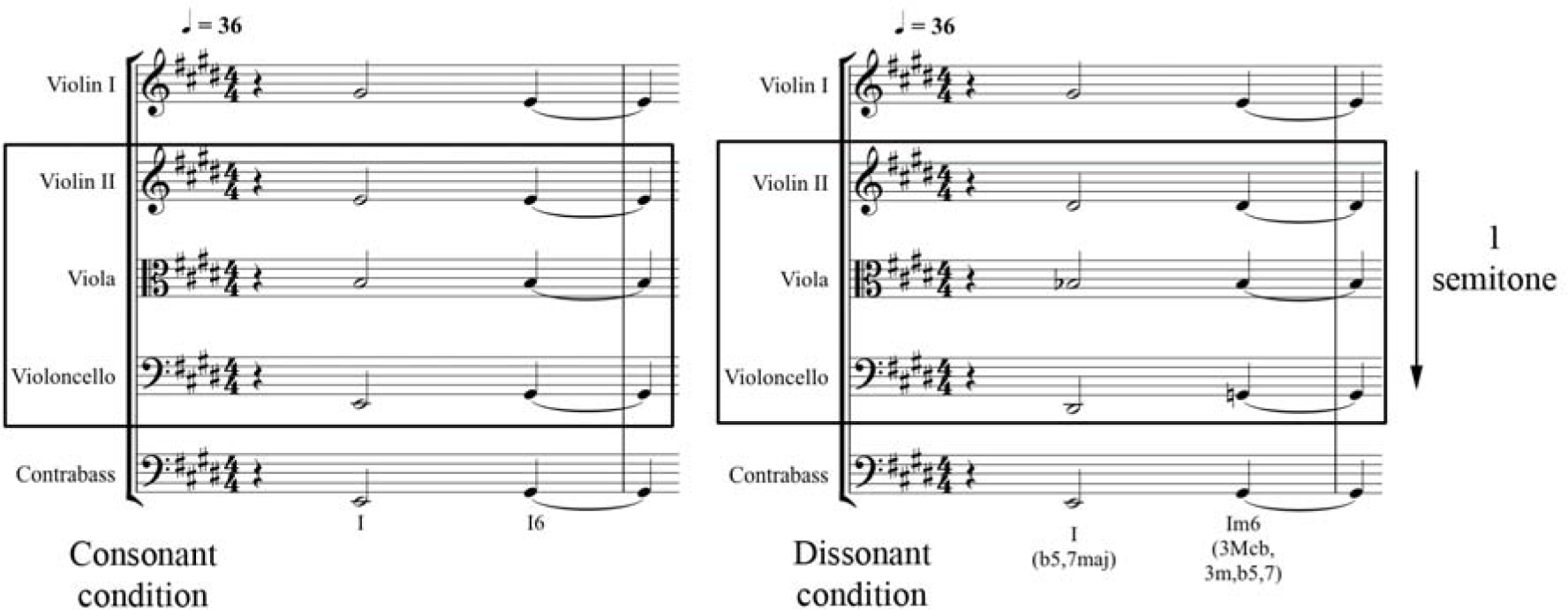
First measure of the consonant (left) and dissonant (right) music conditions. The dissonant condition was obtained by lowering, by a semitone, the second violin, viola and violoncello lines, while keeping the other instruments (first violin and double bass) at their original pitch.

The level of tonal dissonance (Bigand et al., 1996; Bigand & Parncutt, 1999) was the manipulated variable, while other factors that are also known to contribute to the building and release of musical tension such as instrumental timbre, dynamics, rhythm, textural density and melodic contour were strictly controlled (Barthet et al., 2010; Lerdahl & Krumhansl, 2007; Menon, 2002; Paraskeva & McAdams, 1997). The particular experimental transformation used to create the contrasting conditions further allowed the level of consonance/dissonance to be uniform throughout a given category. The scores for the consonant and dissonant conditions were written in Sibelius (version 6.2, Avid, www.sibelius.com), exported as MIDI files (musical instrument digital interface), and played with the same virtual instruments (Strings audio samples from Sibelius 6.2 Library - Sound Essentials). The complete set of materials employed are available to download as supplementary materials. Further information about stimuli construction is available in (Bravo, 2013; Bravo et al., 2019).

Table 3 summarizes interval class content (Forte, 1820; Temperley, 2010) for the first measure of the consonant and dissonant conditions (Figure 5), and portrays the comparative level of consonance/dissonance throughout a given category.

**Table 3.** Interval class content for the first measure of the consonant and dissonant conditions. The model only considers interval classes (i.e., unordered pitch-intervals measured in semitones). Each digit stands for the number of times an interval class appears in the set.

| Interval class | 1 | 2 | 3 | 4 | 5 | 6 |
| --- | --- | --- | --- | --- | --- | --- |
| Consonant condition | 0 | 0 | 3 | 6 | 5 | 0 |
| Dissonant condition | 4 | 2 | 2 | 3 | 6 | 2 |

The consonant condition is primarily governed by collections of intervals considered to be consonant (thirds, fourths and fifths). Whereas, the dissonant condition is characterized by the presence of strongly dissonant intervals (major seconds, minor seconds and tritones). As aforementioned, the music conditions were constructed in the form of variations, with cut-points located at the end of each musical phrase/variation (corresponding to the last chord of each system in Figure 6), resulting in one set of four consonant variations (*consonant condition*: variations C1, C2, C3 and C4) and another set of four dissonant variations (*dissonant condition*: variations D1, D2, D3 and D4). Each variation had a duration of 30 seconds, and consisted of 9 chords presentations (variations 1, 2, 3) or 10 chords (variation 4), totalling 37 chord presentations per music condition. Although the chords belonging to only one music variation per category were found to be sufficient to elicit distinctive valence inferences between conditions at a behavioural level [pre-test: significant differences observed when comparing variations C1 vs. D1; F (1, 27) = 23.637, *p* < 0.001], the experiment was designed to include the complete set of variations per condition, following a strategy similar to that we applied in Bravo et al. (2019); in order to reliably estimate the haemodynamic response function (HRF) and to show detectable differences between conditions in the neuroscientific setting.

Figure 6 (bottom) shows the distribution of pitch-classes for each condition employing the pcdist1 function [MIDI Toolbox for Matlab (Eerola & Toiviainen, 2004)] to calculate the pitch-class distribution of the notematrix (Eerola & Toiviainen, 2004). From the resulting graph, it can be inferred that the consonant condition clearly conveys an E major key content, while the dissonant condition, which spreads through the entire chromatic scale, articulates a much more ambiguous tonal centre.

**Fig. 6.**
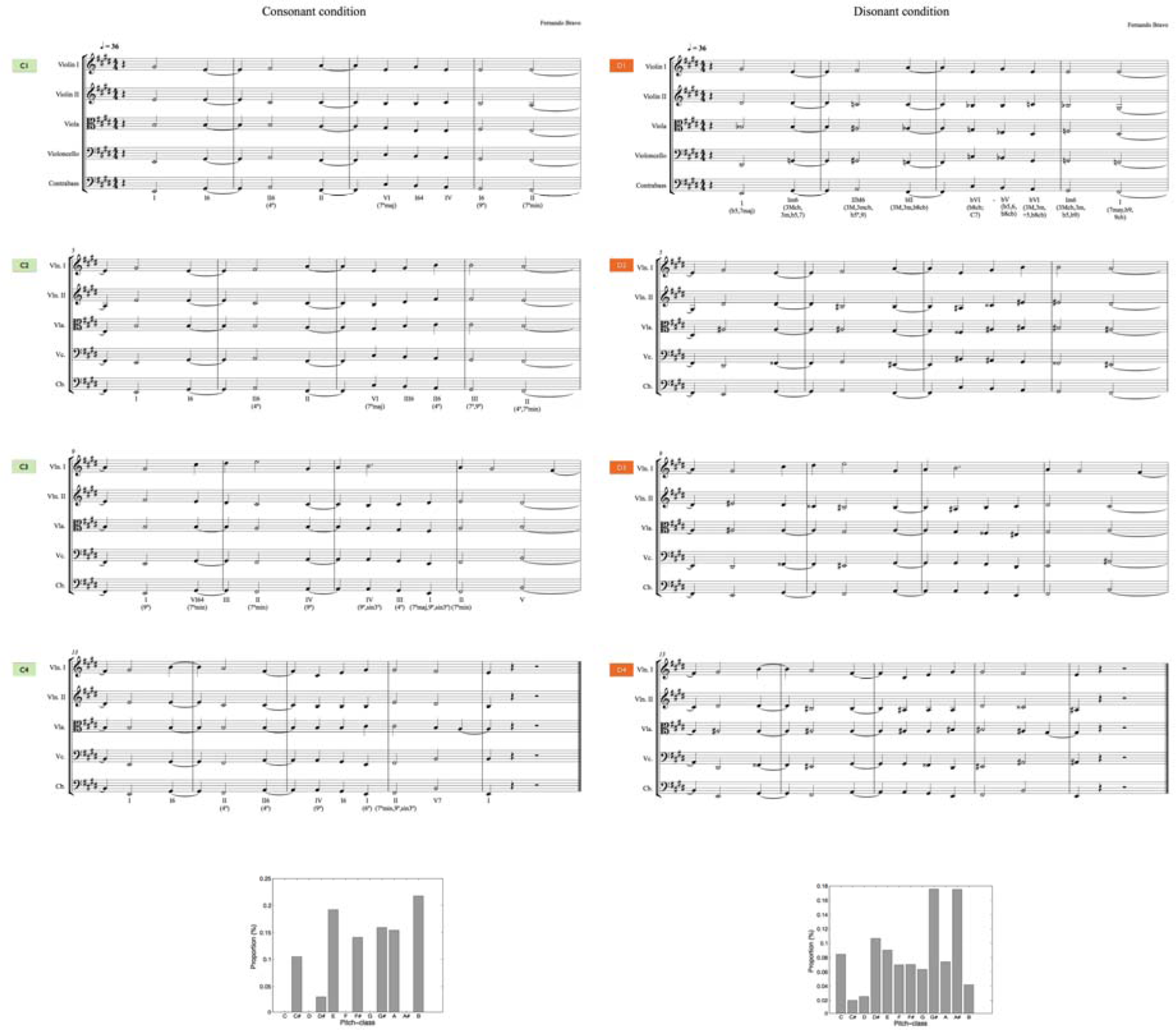
(Top) Complete scores for the consonant and dissonant music conditions. The musical stimuli were written in the form of variations with a duration of 30 seconds each: four consonant variations (*consonant condition*: variations C1, C2, C3 and C4) and four dissonant variations (*dissonant condition*: variations D1, D2, D3 and D4). (Bottom) Key profile for the consonant (left) and dissonant (right) conditions [analysis conducted with MIDI Toolbox (Eerola & Toiviainen, 2004)].

#### Tonalness as a quantifiable predictor for valence inferences to sound

Tonalness has been defined as ‘the degree to which a sonority evokes the sensation of a single pitched tone’ (Parncutt, 1989), in the sense that musical passages with high tonalness evoke the perception of a clear tonal centre (Krumhansl, 2001), whereas sonorities with lower tonalness elicit more unpredictable and equivocal tonal centres. As a component of consonance/dissonance, tonalness has been characterised as the ease with which the ear/brain system can resolve the fundamental, being the easier, the more consonant. Previous studies have consistently shown that the degree of tonalness correlates strongly with the percept of valence (Bravo, 2014; Bravo, Cross, Hawkins, et al., 2017; Bravo, Cross, Stamatakis, et al., 2017). We applied a quantitative probabilistic framework, Temperley’s Bayesian model of tonalness (Temperley, 2002, 2010), to estimate the tonalness level for the two music conditions employed in this experiment. Temperley’s (2010) method relies on the Bayesian ‘structure-and-surface’ approach, and computes tonalness as the overall probability of a pitch-class set occurring in a tonal piece. For musical sequences of a length similar to that used in this study, the approach takes the joint probability of a passage with its most likely analysis, the maximum value of P(structure ∩ surface), as indicative of the overall probability of the passage (Temperley, 2010).

#### The tonalness measure is defined as

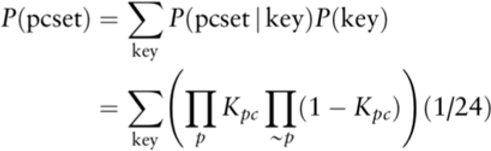

Owing to the uniform manipulation applied to the stimuli in the present experiment, the model predicted a fixed level within the condition. Table 4 shows the two music conditions used in this study, along with their corresponding tonalness values. The fact that the dissonant condition contained multiple chromatic pitch classes meant that there were less facilitated tonal centres, thus resulting in a lower value as assessed by the model.

**Table 4.** Tonalness values for the two sound conditions utilised in the experiment calculated using the Kostka-Payne key-profiles (Kostka, 2003; Kostka et al., 2012).

| Condition | Pitch-class set prime form | Tonalness |
| --- | --- | --- |
| Dissonance | [0 1 3 5 6 8] | 0.00049 |
| Consonance | [0 1 2 3 4 5] | 0.000006 |

The effect of the two music conditions used in this study on emotional valence ratings had been previously established (Table 5). Notably, no significant differences between the two conditions were found either in the arousal or in the potency dimensions (Bravo et al., 2019). In other words, the specific musical structure manipulation employed enabled to target emotional valence and isolate this dimension from arousal and potency.

**Table 5.**
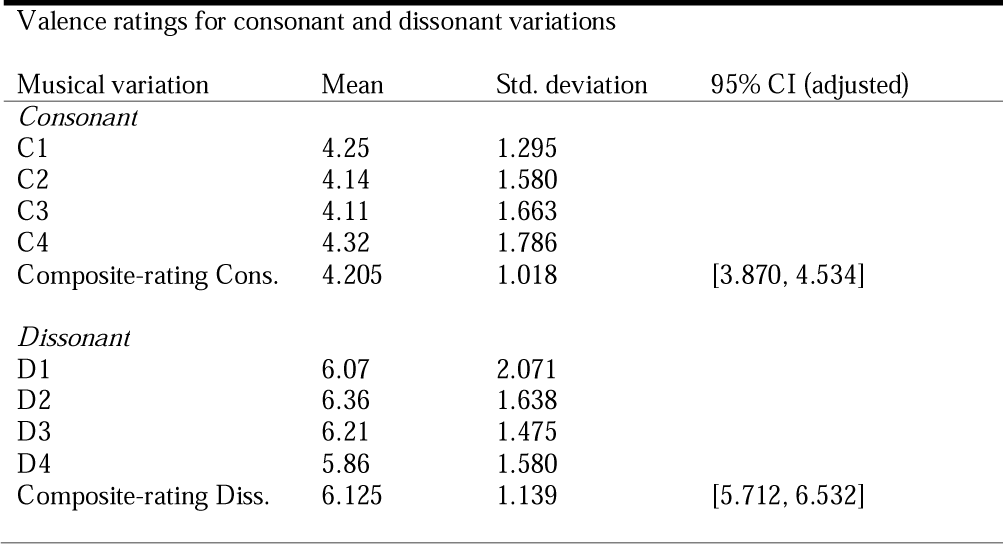
adapted from (Bravo et al., 2019) music listening study]. Individual and composite ratings for consonant and dissonant variations in the valence dimension [mean, standard deviation and, 95% confidence intervals showing adjusted values for repeated measures. Consonant variations are named C1, C2, C3 and C4, dissonant variations are named D1, D2, D3 and D4. [valence: 1 (positive) to 9 (negative)]. Multivariate tests were significant for the valence dimension (there were no significant differences either in the arousal dimension or in the potency dimensions). A paired samples *t* test indicated that the valence composite-rating for all four consonant variations was on average significantly more positive than the valence composite-rating for all four dissonant variations, *t* (27) = 6.87, *p* < 0.001.

| Valence ratings for consonant and dissonant variations |  |  |  |
| --- | --- | --- | --- |
| Musical variation | Mean | Std. deviation | 95% CI (adjusted) |
| *Consonant* |  |  |  |
| C1 | 4.25 | 1.295 |  |
| C2 | 4.14 | 1.580 |  |
| C3 | 4.11 | 1.663 |  |
| C4 | 4.32 | 1.786 |  |
| Composite-rating Cons. | 4.205 | 1.018 | [3.870, 4.534] |
| *Dissonant* |  |  |  |
| D1 | 6.07 | 2.071 |  |
| D2 | 6.36 | 1.638 |  |
| D3 | 6.21 | 1.475 |  |
| D4 | 5.86 | 1.580 |  |
| Composite-rating Diss. | 6.125 | 1.139 | [5.712, 6.532] |

#### Familiarity ratings

Familiarity ratings were measured in (Bravo et al., 2019), which revealed no significant differences between the two music conditions, indicating that the musical materials in both conditions appeared to be similarly unfamiliar to all subjects.

### Procedure

#### Functional Magnetic Resonance Imaging (fMRI) Experiment

Prior to the start of the audio-visual film clip, and based on established methods (Gallagher et al., 2000; Saxe, 2010; Saxe & Kanwisher, 2003; Völlm et al., 2006), an instruction was given to participants designed to engage affective mental state inference processes. Subjects were instructed to “Please, think about the intentions of the main character in the following film clip” (Figure 7). The paradigm, therefore, required to ascribe mental states (e.g., intentions) to an actor on-screen (Van Overwalle, 2009; Van Overwalle et al., 2009). After the instruction, the film clip followed with either music condition 1 (consonance), music condition 2 (dissonance) in randomized order (Figure 7). At the end of the respective clip, a valence inference question was presented (“What type of intentions does the main character have?”), which subjects responded using a 4-point scale (ranging from 1 to 4) with extremes labelled “positive” or “negative” (order counterbalanced).

**Fig. 7.**
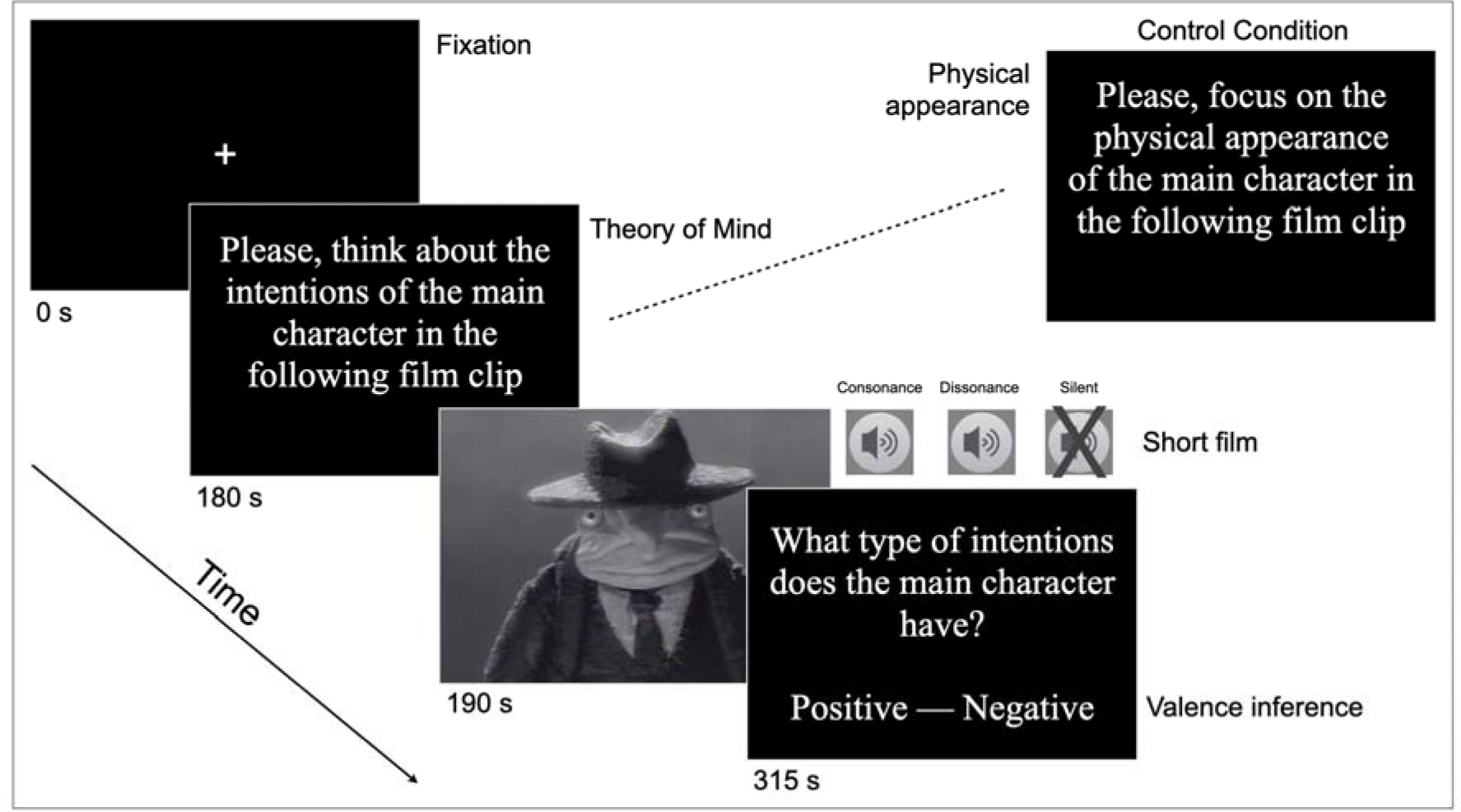
Experimental stimuli and design: After 180 seconds of fixation, an instruction to attend specifically to mental states is given (10 seconds), the film clip [Man with pendulous arms” (Laurent Gorgiard, 1997); duration: 125 seconds] follows with either music condition 1 (consonance) or condition 2 (dissonance) in randomized order. Following the clip participants are asked to rate the valence of the movie character’s intentions (10 seconds) on a scale ranging from one (good intentions) to four (bad intentions). Two control conditions are included; i) visual alone category (i.e., film clip with no soundtrack but same instruction as above) and, ii) film clip with an instruction to describe the “physical” appearance of the character, to control for multimodal sensory processing, working memory and attentional demands of the task.

Two control conditions were included; i) visual alone category (i.e., film clip with no soundtrack) with identical instructions as above to control for basic visual sensory processing, and; ii) the same audio-visual film clips with an instruction to describe the “physical” appearance of the character on-screen (“Please, focus on the physical appearance of the main character in the following film clip”). This additional control condition was aimed at controlling for multimodal sensory processing, working memory and attentional demands of the task, without cueing subjects to attend specifically to mental states (Happé, 1994; Saxe, 2010; Völlm et al., 2006).

The experiment was run in the Neuroimaging Center (NIC) at TU Dresden. Participants were asked to arrive 45 min before the fMRI scanning in order to undertake a preparatory session of 10 minutes in a separate room (contiguous to the scanner room). Subjects were familiarized with the task and trained on the procedure, including the use of the 4-point scale response box system. In the fMRI setup, the audio-visual stimulus was projected onto a screen and presented to the subject via a 45° angled mirror positioned above the participant’s head. The auditory stimuli were delivered via MRI-compatible headphones. Sound pressure levels were measured with a Galaxy Audio CM130 Meter, the output volume was set to 70 dB. Participants were instructed to avoid head movements throughout the functional scan, to keep their eyes on the screen throughout the session and to focus on the presented audio-visual stimuli. Following the scanning session, each subject completed a questionnaire in order to collect subject specific socio-demographic information.

### MRI Data Acquisition

Scanning was performed with a 3.0 T system (General Electric, Signa). Prior to the functional magnetic resonance measurements, high-resolution (1 x 1 x 1 mm) T1-weighted anatomical images were acquired from each participant using three-dimensional fast spoiled gradient-echo (3D-FSPGR) sequence. Continuous Echo Planar Imaging (EPI) with blood oxygenation level-dependent (BOLD) contrast was utilised with a TE of 40ms and a TR of 3000ms. The matrix acquired was 64 x 64 voxels (in plane resolution of 3 mm x 3 mm). Slice thickness was 4 mm with an interslice gap of 0.7 mm (35 slices, whole brain coverage).

### fMRI Data analysis

Data were processed using Statistical Parametric Mapping (SPM), version 12 (www.fil.ion.ucl.ac.uk/spm). Following correction for the temporal difference in acquisition between slices, EPI volumes were realigned and resliced to correct within subject movement. A mean EPI volume was obtained during realignment and the structural MRI was coregistered with that mean volume. The coregistered structural scan was normalized to the Montreal Neurological Institute (MNI) T1 template (Friston et al., 1995). The same deformation parameters obtained from the structural image, were applied to the realigned EPI volumes, which were resampled into MNI-space with isotropic voxels of 3 cubic millimetres. The normalized images were smoothed using a 3D Gaussian kernel and a filter size of 6 mm FWHM. A temporal high-pass filter with a cut-off frequency of 256Hz was applied with the purpose of removing scanner attributable low frequency drifts in the fMRI time series.

An event-related design was employed (Henson, 2007), modelling each of the thirty-seven chord presentations per music category using a canonical haemodynamic response function (Friston et al., 1999) (Model specification timings are available together with the fMRI data in the figshare repository). The design matrix for the first level analysis included the following four regressors: consonance (i.e., movie with consonant music); dissonance (i.e., movie with dissonant music), visual alone (i.e., movie without music); control (i.e., movie with dissonant or consonant music with a control task-physical appearance-). Parameter estimate images were generated. Four contrast images per individual were calculated: dissonance > consonance, consonance > dissonance, consonance > visual alone, dissonance > visual alone, and consonance/dissonance task > consonance/dissonance control. The second level group analysis was carried out using one-sample t-tests. The significant map for the group random effects analysis was thresholded at *p* < 0.001 for voxel-level inference with a cluster-level threshold of *p* < 0.05 corrected for the whole brain volume using FWE (family wise error), which controls for the expected proportion of false-positive clusters.

Whole-brain analyses were performed for all linear contrasts except for the comparison between consonance/dissonance task (ToM) > consonance/dissonance control (physical appearance). The latter contrast was restricted to regions of interest that were defined based on meta-analytic reviews (statistical summaries of empirical findings across studies) that have investigated the neural response to tasks involving theory-of-mind processing (Table 6). All ROIs were defined using anatomical masks of the described areas with WFU PickAtlas Toolbox (Maldjian et al., 2003).

**Table 6.** Regions of interest for fMRI analysis of the contrast between consonance/dissonance task > consonance/dissonance control.

| Region of Interest | Motivation | References |
| --- | --- | --- |
| Anterior cingulate cortex and medial prefrontal cortex | Mental state attribution | Saxe and Kanwisher, 2003; Saxe and Wexler, 2005 |
| Temporo-parietal junction (angular gyrus, supramarginal gyrus and superior temporal gyrus) | Processes requiring intentional attribution and temporary inferences | Saxe, 2006, 2010, Scholz et al. 2009; Carter and Huettel, 2013; Van Overwalle, 2009 |

### Psycho-physiological interactions (PPI) analysis

Psycho-physiological interactions analysis (PPI) was carried out following the approach developed by Friston et al. (1997). Seeds ROI in the left middle occipital gyrus and in the right calcarine cortex were selected on the basis of significantly activated clusters (at threshold level *p* < 0.001 voxel uncorrected, *p* < 0.05 cluster FWE-corrected) from the subtractive analysis for the contrast dissonance > consonance. The group cluster peaks (MNI: −15 −94 8) in the left middle occipital cortex; and (MNI: 12 −94 8) in the right calcarine cortex, were used as point of reference to identify individual subject activation peaks that complied with the following two rules: a) were within a 24 mm radius, and b) were within the boundaries of the corresponding brain area [defined using the WFU pickatlas toolbox: (Lancaster et al., 2000; Maldjian et al., 2003)].

After the identification of the relevant statistical peaks for each subject, spheres were defined around these peaks with a 6 mm radius, which were used as the seed regions of interest (ROIs) for the Psychophysiological Interaction (PPI) analysis. This type of analysis is used to detect target regions for which the covariation of activity between seed and target regions is significantly different between experimental conditions of interest. At the first level, contrasts were calculated for each subject based on the interaction term between the contrast of interest (dissonance > consonance) and each seed ROI’s activity time-course (Friston et al., 1997). For each seed ROI, the contrast images from all subjects were used in voxel-wise one-sample *t*-tests at the second level (at threshold level *p* < 0.001 voxel uncorrected, *p* < 0.05 cluster FWE-corrected) to identify clusters of voxels for which the psychophysiological interaction effect was significant for the group.

### Dynamic causal modelling (DCM) analysis

Effective connectivity analysis was conducted using dynamic causal modelling (DCM) (Friston, 2003). DCM can be applied to networks with as few as two nodes, and some of its most powerful applications have been performed with very simple models (Stephan et al. 2010). In this study DCM analysis was conducted for each subject on two ROIs: the right middle temporal gyrus (rMTG) and the left middle occipital gyrus (lMOG). Coordinates for defined areas are detailed in Table 7. The modulatory effect of interest was the dissonance condition, since this category appeared to strongly modulate the functional interaction between these two areas (as shown in the PPI analysis).

**Table 7.** Regions of interest for dynamic causal modeling (DCM) analysis, describing peak activation MNI coordinates for clusters examined, mask definition and contrasts for VOI extraction (Cl.: cluster, MTG: middle temporal gyrus, MOG: middle occipital gyrus).

| Region of Interest | Cl. Peak | Contrast for Mask | Contrast for VOI extraction | Modulatory effects examined |
| --- | --- | --- | --- | --- |
| Left middle occipital gyrus | -15 -94 8 | Dissonance > Consonance | Dissonance > Consonance | Dissonance |
| Right middle temporal gyrus | 51 -70 -1 | PPI analysis, seed in right PHC<br>(Diss>Cons) | Dissonance > Consonance | Dissonance |

DCM utilises the temporal information contained in fMRI data to estimate and make inferences about the causal relationships of activity patterns between different brain areas, by comparing the evidence for different models of the same data, where each model represents a mechanistic hypothesis about how the data were generated (Friston et al. 2003).

For each subject, three models were defined. Model 1 specified a connection from the rMTG to the lMOG (hypothesized). Model 2 specified a connection from the lMOG to the rMTG. Model 3 specified bidirectional connections between both regions. To explore whether dissonance induced changes in connectivity between brain areas, this condition was included as a modulatory effect allowing it to change any connection in the model. A random effects approach was used since it could not be assumed that the best fitting model structure would be constant across subjects (Stephan et al. 2010). Bayesian model selection (BMS) was utilised to select the optimal model (Stephan et al. 2010). DCM estimates three kinds of coupling parameters for a given model: (i) Direct influences of driving inputs on the neuronal states, (ii) strengths of intrinsic connections that reflect the context-independent coupling between neuronal states in different regions, and (iii) modulatory or bilinear inputs that reflect context-dependent changes (i.e., produced by the experimental conditions) in the coupling between regions. Extrinsic parameters were not estimated in the present analysis, since the process of interest specifically targeted modulatory parameters, which measured changes in effective connectivity induced by dissonance. The parameter estimates describe the speed at which the neural population response changes, which has an exponential decay nature (Stephan et al. 2010). Therefore, parameters are expressed in terms of the rate of change (unit: Hz) of neuronal activity in one area that is associated with activity in another, and can be either positive or negative. A positive parameter means that an increase in activity in one region results in increased rate of change in the activity of another region. Conversely, a negative parameter means that an increase in activity in one region results in a decreased rate of change in the activity of another region. DCM was performed using SPM12 software (http://www.fil.ion.ucl.ac.uk/spm).

## Supporting information

Audiovisual stimuli

## Acknowledgements

We thank P. Heaton, R. Allen, I. Cross, S. Koelsch, F. Franco and M. Rohrmeier for helpful discussions. We thank Christine Ahrends, Katharina Pitt and Aya Keller for help with data collection. Fernando Bravo is funded by a Research Fellowship from the University of Tübingen. This project was made possible through the support from Queens’ College Cambridge (Walker Studentship to F. Bravo), the Andrea von Braun Stiftung, and Zukunftskonzept at TU Dresden (Exzellenzinitiative of the Deutsche Forschungsgemeinschaft). The funding sources had no involvement in the study design or the collection, analysis, and interpretation of the data.

